# Synthesizing Disparate LiDAR and Satellite Datasets through Deep Learning to Generate Wall-to-Wall Regional Forest Inventories

**DOI:** 10.1101/580514

**Authors:** Elias Ayrey, Daniel J. Hayes, John B. Kilbride, Shawn Fraver, John A. Kershaw, Bruce D. Cook, Aaron R. Weiskittel

**Affiliations:** School of Forest Resources, University of Maine, 5755 Nutting Hall, Orono, Maine, 04469-5755, USA; College of Earth, Ocean, and Atmospheric Sciences, Oregon State University, 114 Wilkinson Hall, Corvallis, OR 97331, USA; Faculty of Forestry and Environmental Management, University of New Brunswick, PO Box 4400, 28 Dineen Drive, Fredericton, NB, E3B5A3, Canada; NASA Goddard Space Flight Center, Biospheric Sciences Laboratory, Code 618, Greenbelt, MD, 20771, USA

## Abstract

Light detection and ranging (LiDAR) has become a commonly-used tool for generating remotely-sensed forest inventories. However, LiDAR-derived forest inventories have remained uncommon at a regional scale due to varying parameters between LiDAR datasets, such as pulse density. Here we develop a regional model using a three-dimensional convolutional neural network (CNN), a form of deep learning capable of scanning a LiDAR point cloud as well as coincident satellite data, identifying features useful for predicting forest attributes, and then making a series of predictions. We compare this to the standard modeling approach for making forest predictions from LiDAR data, and find that the CNN outperformed the standard approach by a large margin in many cases. We then apply our model to publicly available data over New England, generating maps of fourteen forest attributes at a 10 m resolution over 85 % of the region. Our estimates of attributes that quantified tree size were most successful. In assessing aboveground biomass for example, we achieved a root mean square error of 36 Mg/ha (44 %). Our county-level mapped estimates of biomass were in good agreement with federal estimates. Estimates of attributes quantifying stem density and percent conifer were moderately successful, with a tendency to underestimate of extreme values and banding in low density LiDAR acquisitions. Estimate of attributes quantifying detailed species groupings were less successful. Ultimately we believe these maps will be useful to forest managers, wildlife ecologists, and climate modelers in the region.

## 1. Introduction

### 1.1 Overview

Over the past two decades light detection and ranging (LiDAR) has become a common tool for developing spatially-explicit forest inventories (Naesset 1997). Measurements of point cloud datasets derived from LiDAR can be used to predict useful forest inventory attributes such as biomass, stem volume, tree count, and species (Means et al. 2000, Jensen et al. 2006, Lim and Treitz 2004). These inventories are useful for a wide range of applications, including assessing carbon stocks (Patenaude et al. 2004), assisting in precision forestry (Woods et al. 2011), and predicting wildlife habitat (Wulder et al. 2008, García-Feced et al. 2011).

Forest inventories are typically developed using the *area*-*based* approach (White et al. 2013), where the forest is segmented into a series of grid cell-areas, ranging from 10 m to 1 ha in size. First, the LiDAR point cloud and the desired forest attribute (e.g. stem density) within each grid cell are each measured. Next, predictive models are then developed relating the field measurements to the LiDAR measurements. Finally, these models can then be applied to every grid cell across a landscape. The resulting maps are referred to as enhanced forest inventories (EFIs).

While LiDAR data are becoming increasingly available to the public, few studies have emphasized mapping whole regions, focusing instead on specific municipalities or individual parcels. One example of regional LiDAR modeling occured in Sweden, which recently completed a project to develop nation-wide forest inventory maps at a 12.5 m resolution (Nilsson et al. 2017). Similar maps have also been generated in Finland (Kangas et al. 2018). In Canada, large portions of Alberta’s forests have had their vegetative functional groups mapped (Guo et al. 2017), and in New Brunswick, a province-wide effort has resulted in near wall-to-wall LiDAR inventories (Dick 2019). We note that each of these examples make use of largely homogenous LiDAR datasets.

### 1.2 The current approach

One common difficulty in generating regional LiDAR inventories is that many regions are comprised of a patchwork of LiDAR datasets acquired with various specifications, and with forest analytics as a secondary objective. This is particularly problematic because the standard approach for measuring a LiDAR point cloud for the development of an EFI model is to take a series of summary statistics describing point heights and vertical distributions. These include measures of the mean, variance, and vertical quantiles, as well as proportions of points that fall above certain height thresholds (McGaughey 2009, Silva et al. 2017). Unfortunately these traditional features suffer from several drawbacks, such as (1) a high degree of autocorrelation, (2) a propensity to change based on LiDAR sampling design (e.g. pulse density), and (3) a propensity to change based on phenology.

Several software suites exist for extracting height features from LiDAR, each producing upwards of fifty features including the heights of every 10^th^ percentile (Silvia et al. 2015). While powerful predictors, many of these measurements are also highly correlated, and so there is a risk of model overfitting without careful model selection (Junttila et al. 2015, Næsset et al. 2005). Unfortunately, many studies make use of every predictor and report on those most important, despite most modelling techniques being unreliable for ranking autocorrelated features (,).

A standard measure for assessing LiDAR quality is pulse density, which refers to the number of laser pulses landing within a given area (pls/m^2^). Pulse density can vary both between and within LiDAR acquisitions, and frequently regional LiDAR collections consist of many acquisitions in which pulse density varies by up to an order of magnitude. Many studies have found that varying pulse density can adversely affect EFI predictions across different LiDAR data sets. For example, in Gobakken and Næsset (2008) noted that the area-based approach was strongly effected by pulse density. Hansen et al. (2015) determined that EFI estimates could be subject to biases if predictions were made on point clouds of different densities than those used to train the model. Other differences in acquisition parameters related to LiDAR sampling design – such as sensor type, pulse frequency, and flight altitude – can also result in different height features being generated over the same area of forest (Næsset 2009, Goodwin et al. 2006). Seasonality also has a major impact on LiDAR height features (Næsset 2005, White et al. 2015, Villikka et al. 2012). In deciduous forests, the presence of leaves can result in LiDAR beams being intercepted higher in the canopy (Ørka et al. 2010). Generally, models developed using one LiDAR point cloud are often not applicable to another, prohibiting regional LiDAR modelling without unusually consistent LiDAR datasets (White et al. 2013).

### 1.3 Deep learning

Here we use deep learning to overcome these obstacles by developing a single model for predicting forest attributes that is applicable across many disparate LiDAR and satellite datasets. Deep learning is a form of machine learning, and primarily refers to artificial neural networks of a sufficient complexity so as to interpret raw data, without a need for human-derived explanatory variables. These differ from simpler neural networks (such as perceptrons) which make estimates using a set of features derived from the data (e.g. height percentiles). Recently, deep learning has proven itself successful at classifying imagery despite varying contextual information, such as light levels and background subject matter (Krizhevski et al. 2012, LeCun et al. 2015). We posit that deep learning will improve EFI modeling by identifying useful spatial features in the LiDAR point cloud without the need for human-derived explanatory variables such as height metrics. These features can be complex shapes and gradients in 3-D space that are less subject to change relative to one another with different acquisition parameters, such as the edges of tree crowns (Ayrey and Hayes 2018).

We implemented a type of spatial deep learning model called a convolutional neural network (CNN). A CNN works by passing a series of moving kernels over spatial data. As the model trains, the weights of those kernels are tuned to identify features useful for predicting the dependent variables (such as the edges of objects). Deep CNNs stack many of these moving windows atop one another, allowing the algorithm to quantify complex features. The three-dimensional CNN developed here uses a volumetric window to quantify a LiDAR point cloud that has been binned into voxel-space. The 3D CNN is thus able to quantify vertical as well as horizontal features, and shapes such as tree crowns, providing a level of complexity not captured by height metrics.

Early CNNs were first developed in the late 1990s and were used to classify hand written digits (Lecun and Bengio 2005). The technique was largely underrepresented in data science until advances in computing power, technique, open-source tools popularized them in 2012 (Krizhevsky et al. 2012). Since then, CNNs of increasing complexity have consistently outperformed models based on feature extraction for computer vision tasks (Szegedy et al. 2015, Tiagman et al. 2014, He et al. 2016). More recently, CNNs have been becoming a popular option for remote sensing problems. There has been much success in using 2D CNNs to classify aerial imagery, hyperspectral, and LiDAR data (Rizaldy et al. 2018, Ghamisi et al. 2017, Castelluccio et al. 2015).

In relation to forestry, some have used segmentation algorithms to isolate individual trees from LiDAR, then used 2D CNNs to classify tree species (Guan et al. 2015, Ko et al. 2018). Work is also being done using 2D CNNs to identify individual tree crowns from high resolution imagery (Weinstein et al. 2019, Li et al. 2016). Progress has also been made in adapting CNNs to scan LiDAR point clouds and three dimensional space. Similar CNNs that make use of voxels to quantify point clouds have been used to identify household objects and malignancies in medical scans (Qi et al. 2016, Maturana and Scherer 2015). In 2017 Qi et al. introduced PointNet, designed to interpret LiDAR data without voxels, although this technique does not make full use of the spatial relationship between neighboring points. In a remote sensing context, Maturana and Scherer (2015) used a 3D CNN to identify helicopter landing zones in LiDAR. Most similar to this study, Ayrey and Hayes (2018) tested a variety of CNN architectures to interpret LiDAR for the estimation of forest attributes.

### 1.4 Objectives

The primary objective of this study is to develop a regional EFI over the Acadian/New England Forest region, with a total of 85 % coverage of the New England states using publically-available LiDAR. We overcome several challenges faced by having disparate LiDAR datasets through the use of deep learning, and we incorporate other remote sensing products such as spectral data, phenology, and disturbance history to improve model accuracy. A secondary objective is to assess the value of deep learning, and compare it to standard approaches for LiDAR modelling. A final objective is to compare our mapped estimates of biomass, percent conifer, and tree count to estimates derived via the design-based U.S. national forest inventory program. We will compare estimates made at the plot-level and at the county-level using both inventory techniques, and identify any obvious shortcomings. The end result will be a series of near wall-to-wall mapped forest inventory estimates of the region, with an accurate assessment of error across space and forest type. This will provide forest managers, ecologists, and climate modelers in the region with an unprecedented amount of detailed information about the forest.

## 2. Materials and Methods

### 2.1 Forest attributes

Our goal was to estimate many forest attributes which may be useful to ecologists, forest managers, and modelers. The complete list is found in Table 1. For brevity, at points throughout this manuscript we will highlight only the results of the aboveground biomass (BIOAG), percent conifer (PC), and tree count (TC) estimations. All other attributes can also be considered measurements of tree size, density, or species; and are often represented by these three.

**Table 1.** A complete list of forest attributes estimated in this study. Note that all estimates were made exclusively on trees greater than 10 cm in diameter.

| Forest Attribute | Units | Description |
| --- | --- | --- |
| Aboveground Biomass (BIOAG) | kg | Aboveground biomass as calculated by the USFS's FIA component ratio method. |
| Total Biomass | kg | Total woody biomass as calculated by the USFS's FIA component ratio method. |
| Basal Area | m<sup>2</sup> | Basal area at breast height. |
| Mean Tree Height | m | Mean height of the trees' apices. Not a measure of mean overall canopy height. |
| Quadratic Mean Diameter | cm | Quadratic mean of diameter at breast height. |
| Volume | m<sup>3</sup> | Total inner bark volume of each tree's bole. |
| Volume, Merchantable | m<sup>3</sup> | Total merchantable inner bark volume of each tree's bole; starting at 10 cm above ground and ending at a height of 10 cm in diameter. |
| Tree Count (TC) | # | Total number of trees. |
| Percent conifer stems (PC) | % | Percentage of conifer stems. |
| Percent spruce/fir | % | Percentage of spruce or fir species stems. |
| Percent *pinus strobus* | % | Percentage of white pine (*Pinus strobus*) stems. |
| Volume of deciduous | m<sup>3</sup> | Total inner bark volume of deciduous tree boles. |
| Volume of spruce/fir | m<sup>3</sup> | Total inner bark volume of spruce or fir tree boles. |
| Volume of *pinus strobus* | m<sup>3</sup> | Total inner bark volume of white pine (*Pinus strobus*) tree boles. |

### 2.2 Training data

For most applications deep learning requires very large datasets (Russakovsky et al. 2015). To meet this requirement we combined thirteen different forest inventories collected over thirty two sites (Appendix Table A1). Within each inventory all trees greater than 10 cm in diameter were measured spatially with species and diameter recorded. In several inventories, tree heights were only measured on only a subset of trees, so species-specific non-linear height-diameter models were generated using a Chapman-Richards model form to impute tree height using the existing height measurements at each site. Some inventories were also measured up to ten years prior to LiDAR acquisition. In instances in which the temporal discrepancy between LiDAR and field data exceeded two years, trees were projected forward in time using the Forest Vegetation Simulator’s Acadian Variant (Weiskittel et al. 2012).

Volume was estimated using regional taper equations (Li et al. 2012). A 10 cm cutoff was used for merchantable volume. Biomass was estimated using the component ratio method developed for the US Forest Inventory and Analysis (FIA) program (Woodall et al. 2011).

Each of the stem-mapped inventories were aligned visually with the LiDAR to correct for GPS error in plot location, and then segmented into 10×10 m grid cell plots. We selected this resolution in order to amplify the amount of unique plots available, while retaining plots large enough to contain several whole tree crowns. While they provide highly spatially-explicit models, this plot size is small compared to those used in other studies, and so can be prone to edge effects.

We accounted for edge effects by using regional diameter-to-crown width equations to project each tree’s crown in space (Russell and Weiskittel 2011). Tree level basal area, biomass, and volume allometry were then multiplied by the proportion that each tree’s crown overlapped the plot. Trees were therefore treated as areas containing biomass, rather than points which could lie one or another side of a plot boundary (,). This mimics the method by which the remote sensing instrument measures the trees, as LiDAR or imagery has no means of measuring the precise location of a tree’s stem. Figure 1 demonstrates this correction. Preliminary testing using a subset of the data indicated that this correction greatly improved model performance.

**Figure 1.**
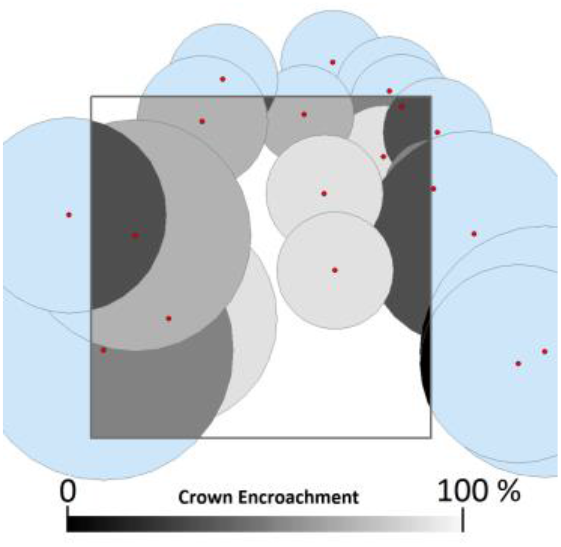
The percent of crown overlap of each tree in and around the plot was used as a modifier on that tree’s basal area, biomass, and volume. This allowed for the development of models which more closely reflected what was visible to the remote sensing systems, while remaining unbiased across multiple cells.

We also augmented the sample size of our training data by allowing plots to overlap one another by a maximum of 25 %, and by including multiple LiDAR acquisitions of the same plot in the training dataset. The precise configuration of LiDAR returns will always vary between acquisitions. Similar augmentation techniques, such as transforming or rotating input images multiple times, and using adjacent still frames of videos have been successfully used in deep learning for many years (). Lastly, we included 500 plots with no trees on them to allow for better predictions in low-vegetation environments, which were sampled randomly across Northern New England using the 2011 National Land Cover Database (). The associated LiDAR point clouds were then manually checked for trees and discarded if trees were found.

Ultimately we assembled 24,606 plots for model training and preliminary evaluation. Of these, we randomly withheld 4,000 plots, including 1,000 for model validation (to determine the optimal stopping point during deep learning training), and 3,000 for model testing (used for model comparison and selection). Augmented plots related to withheld plots were removed from the training dataset, thus the final training set was comprised of 17,432 plots.

### 2.3 Remote Sensing Data

#### 2.3.1. LiDAR

We aggregated 49 public LiDAR datasets from across the region, combining acquisitions of varying pulse density and seasonality. Although much of the pubic LiDAR in the US Northeast is flown leaf-off, we chose to develop models capable of functioning in either state to allow for potential future integration with leaf-on Canadian Maritime data. The training data ultimately consisted of a 53 to 47 % split between leaf-off and leaf-on respectively.

The majority of the LiDAR used for this study was funded and hosted by the US Geological Survey’s national 3D Elevation Program (3DEP). These data were captured in leaf-off conditions between 2006 and 2018 at resolutions ranging from 0.5 to 10 pls/m^2^. We also incorporated LiDAR data acquired by NASA Goddard’s LiDAR Hyperspectral and Thermal Imager (G-LIHT) as well as the National Ecological Observatory Network (NEON). These data were acquired in leaf-on conditions over several of our training sites with pulse densities ranging from 5 to 16 pls/m^2^. Finally, we incorporated several private LiDAR datasets for training, including one each over the Penobscot Experimental Forest, Baxter State Park in Maine, and Noonan Research Forest in New Brunswick. Each of these had an average pulse density of 5-6 pls/m^2^, where the first two were leaf-off and the second leaf-on. Both pulse density and seasonality were included as model predictors.

#### 2.3.2 Satellite variables

We also chose to include satellite derived spectral indices, disturbance metrics, and phenology data in our models. Each of these are spatially contiguous across our study area, and have proven useful for predicting forest attributes (Zheng et al. 2004, Pflugmacher et al. 2012, Clerici et al. 2012). All satellite data processing was done in Google Earth Engine (Gorelick et al. 2017).

Using Sentinel-2b data, we generated maps of six spectral vegetation indices: Normalized Burn Ratio (NBR, Key and Benson 2999), Normalized Difference Vegetation Index (NDVI, Rouse et al. 1973), Normalized Difference Moisture Index (NDMI, Cibula et al. 1992), Red-Edge Chlorophyll Index (RECI, Gitelson at al. 2006), Greenness Index (GI, Hunt et al. 2011) and Triangular Chlorophyll Index (TCI, Hunt et al. 2011). Landsat-8 imagery was used generate three tassel-cap indices (brightness, greenness, and wetness, Crist 1985). This imagery was acquired between 2015 and 2017, imagery from between the 150^th^ and 244^th^ Julian days were used. All images were cloud-masked, then a single median composite was developed for the study area. Resolutions greater than 10 m were resampled to match our plot size.

We also incorporated disturbance history using Landsat 5-8 and the LandTrendr disturbance detection algorithm (Kennedy et al. 2010). LandTrendr works by fitting a maximum of seven linear segments to the yearly medians of a spectral band within each Landsat pixel. Vertices identify dramatic changes in the spectral characteristics of that pixel in time, and often correspond to disturbances. We ran LandTrendr over the greenness and wetness tassel cap indices, and NBR. Instances in which LandTrendr identified a vertex in two or more of the three bands, within two years of one another were retained as disturbances. We then condensed these data into the year of last disturbance and the magnitude of that disturbance (as a percent of vegetation index change).

Finally, we estimated growing season length across our study area using Moderate Resolution Imaging Spectroradiometer (MODIS) data with a resolution of 500 m. This was derived by subtracting the mean dates of greenness onset and senescence, and has been demonstrated to correlate well with site quality, and so may aid models which primarily use height to infer tree size (Xhang et al 2006).

### 2.4 Deep learning modelling

#### 2.4.1 Data preparation

We first prepared the LiDAR data to be scanned by the 3D-CNN by converting it from a point cloud with each data point representing an X,Y, and Z value, to volumetric pixels (voxels). A height-normalized point cloud was voxelized by segmenting each 10 × 10 × 35 m space (representing a grid cell) into 40 × 40 × 105 bins, and then tallying the number of LiDAR points within each bin. Thus, each voxel represented a space of 25 × 25 × 35 cm on our plot. We used vertically rectangular voxels to reduce dimensionality and prevent horizontal features from being lost. Voxel size was determined through qualitative testing of several size configurations using a reduced model form. Ultimately the voxel data took the form of a three-dimensional tensor, over which the kernels of a CNN could be passed. Although it has been noted that CNNs frequently perform better and train faster when applied to standardized data (Krizhevsky et al. 2012), we tried several standardization techniques (z-score, prescience/absence, ect…), and found no such improvement.

#### 2.4.2 Model architecture

Our deep learning model architecture was based loosely on Google’s Inception-V3 (Szegedy et al. 2016), which was determined by Ayrey and Hayes (2018) to be better suited than several other commonly-used CNN architectures. The underlying model-form was converted to interpret 3D data, and care was taken to maintain similar proportional dimensionality to the original model (designed to interpret images with a resolution of 224 × 224). The full model architecture is visualized in Figure 2.

**Figure 2.**
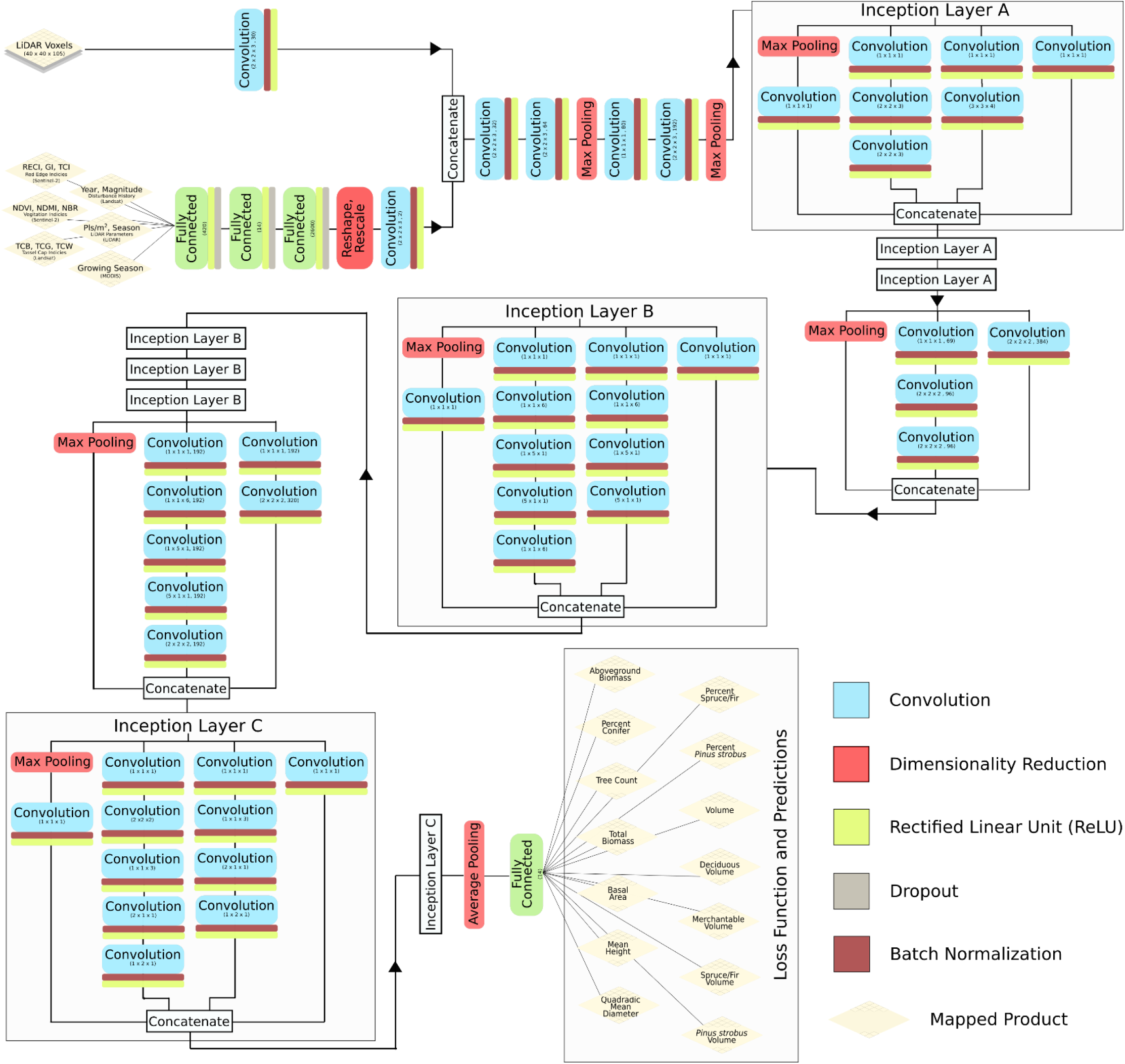
The full architecture of the Inception-VS convolutional neural network used to predict forest attributes from LiDAR and satellite data.

Inception-V3 consists of a series of preliminary convolution and pooling layers, followed by inception layers. Inception layers consist of a number of convolutions of varying sizes which are passed over the incoming data, each designed to pick up on different features, which are then concatenated. Inception-V3 consists of nine inception layers back to back, with intermittent pooling to reduce dimensionality. The final inception layer is fed into a fully connected layer for a classification or regression prediction. Each convolution was followed by a ReLU activation layer and batch normalization.

The model was first trained to estimate only BIOAG using LiDAR. We used a process called transfer learning to initialize the weights of a more complex model using the weights from the simpler one, which was designed to simultaneously predict all 14 of our forest attributes. Each forest attribute was standardized using z-scores so that their values fell upon the same range. A single loss function was then used to optimize the model to predict all forest variables (Equation 1). In which, the mean of the squared error of the *k* standardized forest attributes is summarized to a batch level mean of *n* training samples.

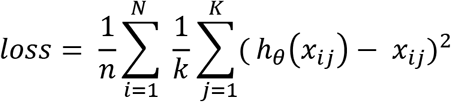

We included the satellite data as side-channel information by first developing a multi-layer perceptron to estimate BIOAG directly from the satellite variables. We took the weights from this model and used them to initialize a subcomponent of the larger model, which produced a 40 × 40 tensor that was then concatenated onto the LiDAR voxels (Zhou et al. 2017).

#### 2.4.3 Training

Deep learning models were developed in Python using Google’s Tensorflow version 1.4 (Abadi et al. 2016). The model training process took place in three stages using transfer learning to build upon each stage (1) A BIOAG model using only LiDAR, (2) A model predicting all fourteen attributes using only LiDAR, and (3) a model predicting all fourteen forest attributes using LiDAR and satellite metrics. Training time for the first stage was approximately five days using an NVIDIA Tesla k80, the following stages were trained more rapidly. This lengthy multi-step training process made cross-fold validation highly impractical.

### 2.5 Traditional modelling

Deep learning models were compared to models trained using the standard suite of LiDAR height metrics, derived using the Rlidar package (Silva et al. 2015). This produces a series of summary statistics of the LiDAR return’s heights and proportions above certain height thresholds. We discarded metrics which made use of LiDAR intensity and return counts, as these could not be normalized between the different LiDAR acquisitions. We filtered out points lower than 0.5 m above ground, and used a 2 m threshold for many of the proportional metrics. Other studies in the region have used similar cutoffs (Hayashi et al. 2014). We also included the aforementioned satellite-derived metrics, as well as pulse density, and seasonality. In total 41 covariates were derived from the LiDAR and satellite data.

Random forest imputation in regression mode was used to model each of the forest attributes (Breiman 2001). Other studies conducted on subsets of our dataset have demonstrated that this modelling technique outperforms linear mixed-effects modelling (Ayrey and Hayes 2018, Hayashi et al. 2015). We used a process called Variable Selection Using Random Forest to eliminate unimportant predictors (Genuer et al. 2015). New models were then developed using 2000 decision trees and one-third variable selection at each node-split. These hyper-parameters were fine-tuned using a subset of the data.

The random forest models were trained and validated using the withheld test plots. Although accuracy can be assessed using out-of-bag sampling, we used the same validation scheme as the deep learning models due to data augmentation and consistency.

### 2.6 Validation

The training, validation, and testing data derived from the thirteen individual forest inventories are likely not fully representative of the landscape, leading to problems with spatial-autocorrelation at the regional scale. We therefore performed two phases of validation. The first phase of validation made use of the 4,000 withheld plots. This was used for model comparisons between deep learning and traditional modelling, and to settle on the final model form.

The second phase of validation made use of an independent dataset, and was used to assess true model performance across the landscape. For this we used the United States Forest Service’s FIA national inventory plots. These consisted of approximately 7,500 stratified-random plots with a nested sampling design within the states of Connecticut, Maine, Massachusetts, New Hampshire, Rhode Island, and Vermont. We used these to determine the maps’ error and bias, assess spatial-autocorrelation across the landscape, and compare our inventory estimates to FIA county-level estimates. We assessed error in Connecticut and Rhode Island separately as their forests increasingly represent a Mid-Atlantic forest type not fully represented by the training data. We removed buildings from our maps using a building mask of the United States developed by Microsoft’s Bing Maps Team using high resolution imagery (Bing Maps Team 2018).

The FIA plots consist of a nested plot design that includes four 7.3 m radius subplots placed 36.6 m away from one another. The subplots have an area of 168 m^2^, while the entire FIA plot taken as an aggregate has an area of 672 m^2^. The individual subplot measurements were more affected by errors in plot location, as these were more subject to intra-canopy variability. A preliminary finding that the center plots (on which the GPS point is taken) produced a lower error than the subplots reinforced this conclusion. The aggregate plot-level measurements were less prone to location errors, but did not necessarily represent the entire range of variability that one would expect in a 10 m grid cell. It is important for validation plots to have roughly the same area as the grid cells being validated so that each have a similar range in values. We therefore assessed accuracy at a subplot- and plot-level, but used the plot-level errors to perform additional analyses. This is roughly equivalent to assessing error were the map resampled to a 20 m resolution.

We decided not to use FIA plots for model training for several reasons: (1) The FIA nested plot design was not compatible with our edge correction technique. (2) FIA plot locations are imprecise and recorded with a consumer-grade GPS, and frequently have an error greater than the size of our cells. (3) FIA plot location data is not publically available and is subject to considerable restrictions, less so if only used for extracting error.

**Table 1.**
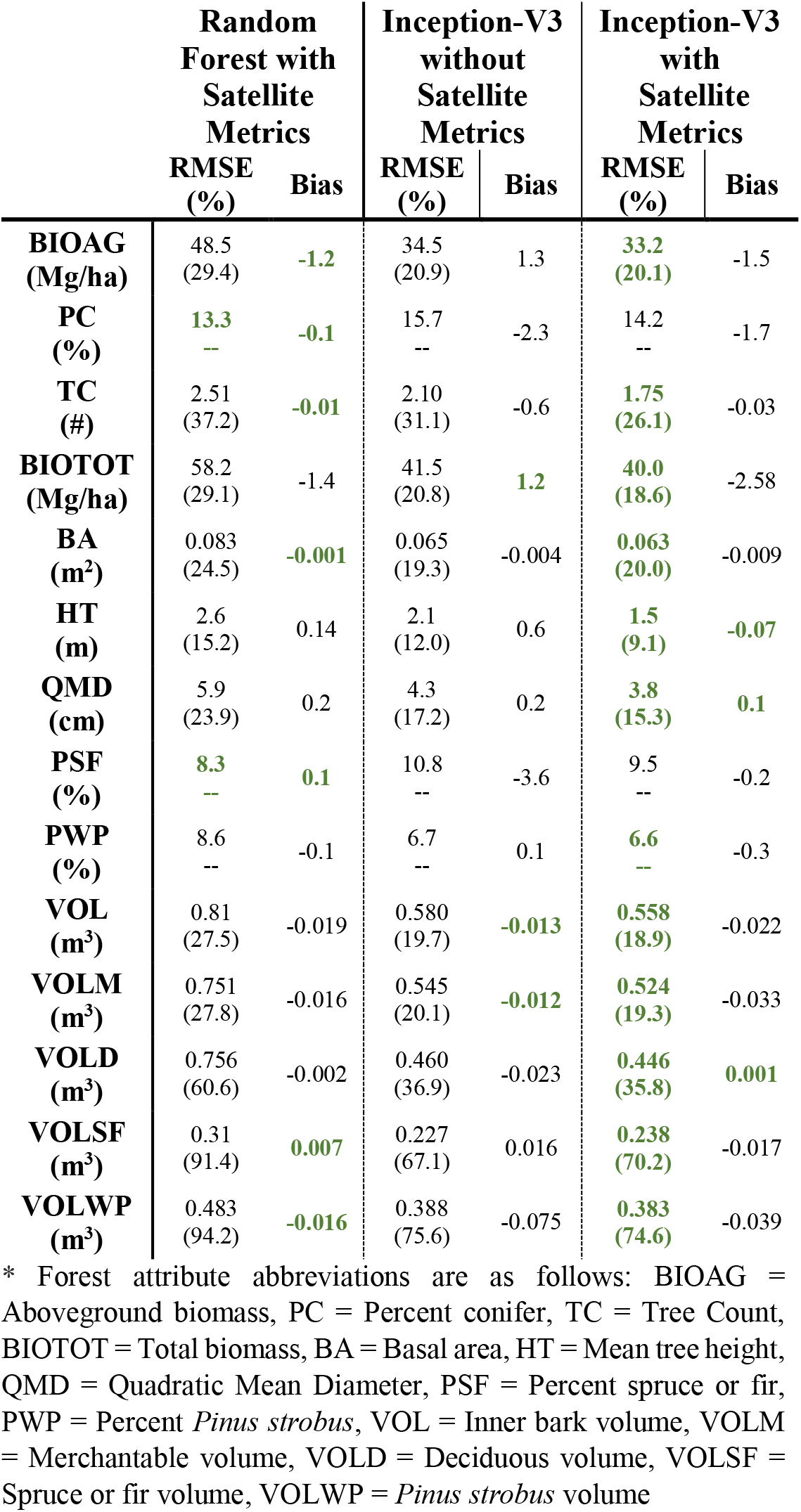
Results in terms of RMSE, RMSE as a percent of mean (%), and bias of three models. Random forest models trained using LiDAR height and satellite metrics, an Inception-V3 model trained using only the LiDAR point cloud, and an Inception-V3 model trained using the LiDAR point cloud and satellite metrics. Highlighted in green are the best values achieved.

|  | Random Forest with Satellite Metrics |  | Inception-V3 without Satellite Metrics |  | Inception-V3 with Satellite Metrics |  |
| --- | --- | --- | --- | --- | --- | --- |
|  | RMSE (%) | Bias | RMSE (%) | Bias | RMSE (%) | Bias |
| **BIOAG (Mg/ha)** | 48.5 (29.4) | **-1.2** | 34.5 (20.9) | 1.3 | **33.2 (20.1)** | -1.5 |
| **PC (%)** | **13.3**<br>-- | **-0.1** | 15.7<br>-- | -2.3 | 14.2<br>-- | -1.7 |
| **TC (#)** | 2.51 (37.2) | **-0.01** | 2.10 (31.1) | -0.6 | **1.75 (26.1)** | -0.03 |
| **BIOTOT (Mg/ha)** | 58.2 (29.1) | -1.4 | 41.5 (20.8) | **1.2** | **40.0 (18.6)** | -2.58 |
| **BA (m<sup>2</sup>)** | 0.083 (24.5) | **-0.001** | 0.065 (19.3) | -0.004 | **0.063 (20.0)** | -0.009 |
| **HT (m)** | 2.6 (15.2) | 0.14 | 2.1 (12.0) | 0.6 | **1.5 (9.1)** | **-0.07** |
| **QMD (cm)** | 5.9 (23.9) | 0.2 | 4.3 (17.2) | 0.2 | **3.8 (15.3)** | **0.1** |
| **PSF (%)** | **8.3**<br>-- | **0.1** | 10.8<br>-- | -3.6 | 9.5<br>-- | -0.2 |
| **PWP (%)** | 8.6<br>-- | -0.1 | 6.7<br>-- | 0.1 | **6.6**<br>-- | -0.3 |
| **VOL (m<sup>3</sup>)** | 0.81 (27.5) | -0.019 | 0.580 (19.7) | **-0.013** | **0.558 (18.9)** | -0.022 |
| **VOLM (m<sup>3</sup>)** | 0.751 (27.8) | -0.016 | 0.545 (20.1) | **-0.012** | **0.524 (19.3)** | -0.033 |
| **VOLD (m<sup>3</sup>)** | 0.756 (60.6) | -0.002 | 0.460 (36.9) | -0.023 | **0.446 (35.8)** | **0.001** |
| **VOLSF (m<sup>3</sup>)** | 0.31 (91.4) | **0.007** | 0.227 (67.1) | 0.016 | **0.238 (70.2)** | -0.017 |
| **VOLWP (m<sup>3</sup>)** | 0.483 (94.2) | **-0.016** | 0.388 (75.6) | -0.075 | **0.383 (74.6)** | -0.039 |

\* Forest attribute abbreviations are as follows: BIOAG = Aboveground biomass, PC = Percent conifer, TC = Tree Count, BIOTOT = Total biomass, BA = Basal area, HT = Mean tree height, QMD = Quadratic Mean Diameter, PSF = Percent spruce or fir, PWP = Percent *Pinus strobus*, VOL = Inner bark volume, VOLM = Merchantable volume, VOLD = Deciduous volume, VOLSF = Spruce or fir volume, VOLWP = *Pinus strobus* volume

## 3. Results

### 3.1 Phase one validation and model comparison

The first validation phase made use of withheld plots to compare fourteen random forest models using height and satellite metrics to two Inception-V3 CNNs. The first CNN made use of only the LiDAR point cloud, while the second made use of the LiDAR point cloud and satellite metrics. Results in terms of RMSE and bias are displayed in Table 1. In terms of RMSE, both CNNs outperformed random forest in predicting 12/14 forest attributes (the two exceptions being for predictions of percent conifer and percent spruce fir). In terms of absolute bias, random forest outperformed both CNNs in 7/14 metrics. It should be noted that in many comparisons of bias, the absolute difference between models was negligible.

The comparison between CNNs with and without satellite metrics illustrated that the satellite metrics are contributing to an improvement in the final model’s performance. In terms of RMSE, the CNN with satellite metrics outperformed the one without 100 % of the time. The CNN without satellite metrics had less absolute bias in predicting 3/14 metrics. Despite this, the proportional improvement in terms of both RMSE and bias after the addition of satellite metrics was relatively small. Additional testing of the random forest model with and without satellite metrics indicated a greater relative improvement in predictive power.

### 3.2 Phase two validation

In the second phase of validation each of the mapped forest attributes were validated across Northern New England using FIA plots. Table 2 displays the results of this validation in terms of RMSE, RMSE as a percent of mean (nRMSE), and bias at both the subplot- and plot-level. We did not assess random forest performance in this phase because we did not apply these models at a regional level; such an effort would have been computationally costly in the extreme and was unnecessary given our model selection process in the first phase. Model error at the subplot-level was considerably higher than error at a plot level with each forest attribute. This is to be expected given that smaller areas are more likely to contain extreme values, and the subplot values are more likely to be affected by GPS inaccuracy. The opposite trend could be seen in bias, with 10/14 attributes exhibiting greater bias at the plot-level.

**Table 2.**
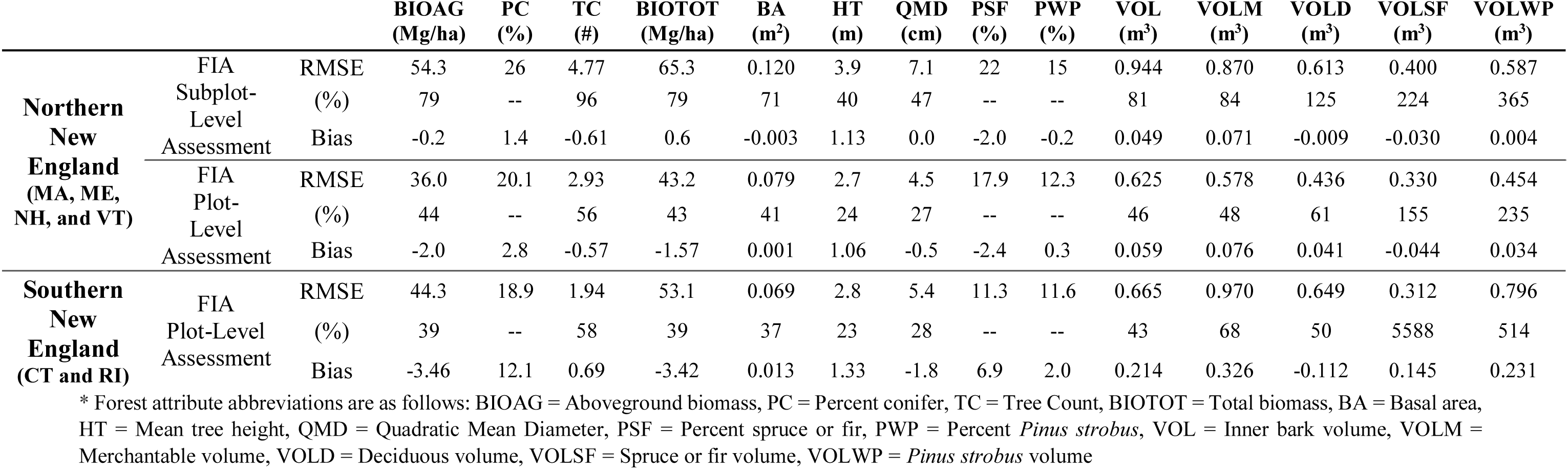
Results in terms of RMSE, RMSE as a percent of mean (%), and bias of the Inception-V3 model using FIA plot data for validation. Assessments were made using the FIA subplots (roughly corresponding to 10 m cell validation) and the FIA plots taken as an aggregate (roughly corresponding to 20 m accuracy).

Results of the second phase of validation indicated that two phases were in fact necessary to obtain a more representative assessment of regional model performance. Plot level RMSE was worse than the RMSE obtained from the first phase of validation in 13/14 forest attributes, indicating that the withheld plots likely did not represent landscape heterogeneity. In some forest attributes this difference was relatively minor. Performance of the tree count, mean height, and species estimates were notably worse in the second phase of validation. The error of each of these values increased between 41-90 % from that observed in the first phase of validation.

Overall, the error and biases of attributes representing tree size were considerably lower than those representing species or stem density. Aboveground biomass, total biomass, basal area, mean tree height, QMD, and volume all had a plot-level nRMSE of less than 50 % in Northern New England. In contrast, tree count had a nRMSE of 56 %. Model performance was worst in volume estimates of species groups. Estimates of the deciduous volume had a relatively high nRMSE, 61 %. Estimates of spruce/fir, and *pinus strobus* volume both had nRMSEs above 150 %, and could generally be considered not useful. We did not assess nRMSE of the percent species group estimates. The RMSE and biases of percent spruce/fir and percent *pinus strobus* were lower than that of percent conifer, but this is likely because their average values are smaller. Qualitatively, the maps of percent conifer appeared better, with the others suffering from more banding and local biases.

We also assessed model performance of BIOAG, PC, and TC in Northern New England spatially and by plotting their predicted verses observed values. Figure 2 illustrates the plot-level residuals of each of the three forest attributes. We note that the BIOAG residuals appear to be fairly evenly distributed across the landscape, with consistent model biases not immediately apparent. Positive PC residuals appear to be clustered mostly in Eastern Maine, where greater number of conifers are likely to be found, indicating that the model is underestimating in areas of proportionally higher conifers. Likewise, negative residuals in Vermont are an indication that the model is underestimating in areas with proportionally fewer conifers. Tree count followed a similar trend, greater residuals were encountered in more northern areas, which correspond to both greater stem density and more industrial forests.

**Figure 2.**
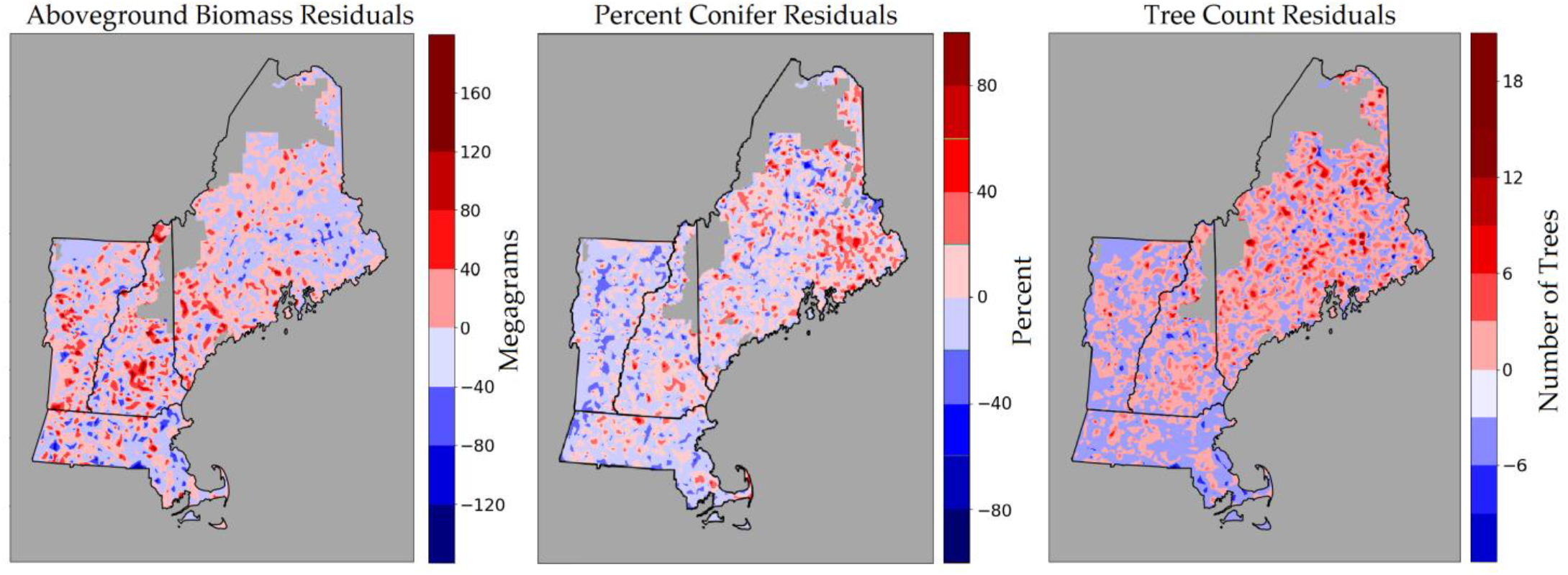
FIA plot-level residuals are plotted for three of mapped forest attributes. Red areas denote model underestimation, blue areas denote model overestimation.

These trends can also be observed in the predicted versus observed plots (Figure 3). Biomass residuals fall relatively tightly along the 1:1 line. Little to no attenuation is observed at higher biomass values. Percent conifer residuals seemed to indicate a tendency to overestimate in low conifer environments, and underestimate in high conifer environments. Finally, tree count residuals appeared to follow the 1:1 line in low-medium density conditions, but often severely underestimated tree count in high-density conditions.

**Figure 3.**
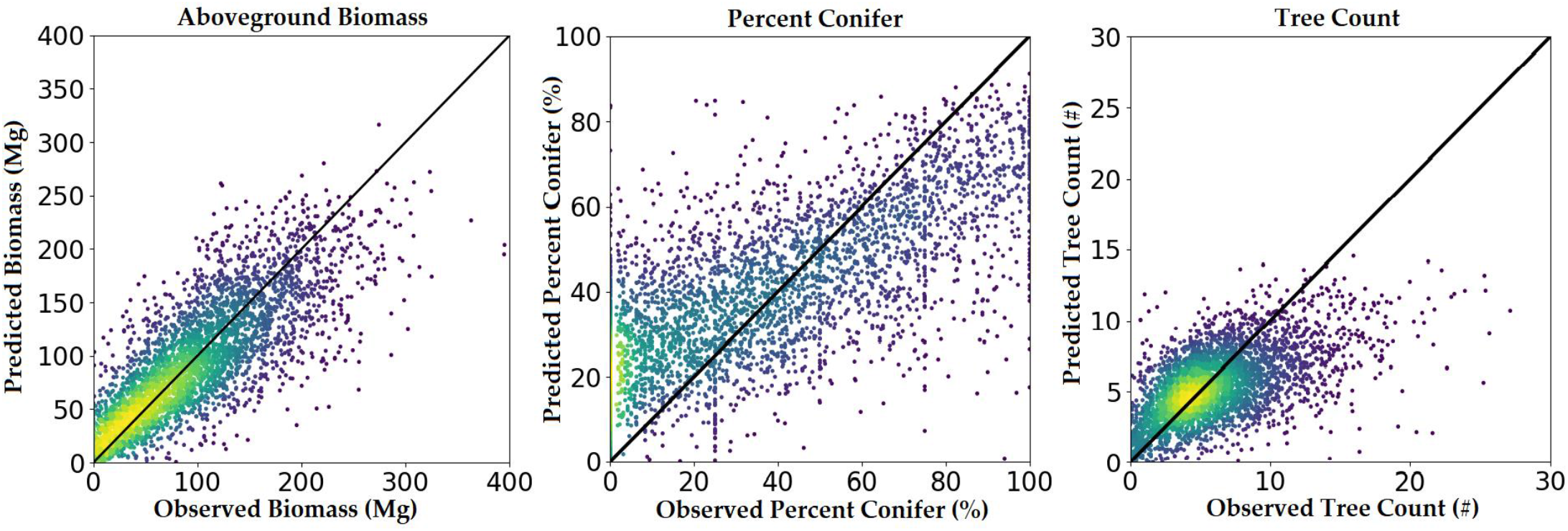
Predicted versus observed plots using FIA plot-level validation.

We assessed model performance across different LiDAR datasets by plotting the Northern New England plot-level bias as a function of pulse density (Figure 4). We used lowess regression to fit a moving trendline to the data using 75 % of observations to smooth the line at each value. No biases stemming from pulse density were apparent in this visualization, with the trend lines for biomass and tree count consistently near-zero, and the trend line for percent conifer showing a slight positive bias, but had no apparent trend with pulse density. Nevertheless, banding was visible on the maps of percent conifer and tree count in regions of very low pulse density. The bands appeared to follow trends in average scan angle along each flight line, and so may have been a function of pulse density and scan angle combined. Unfortunately, we did not map mean scan angle across the landscape a-priori as we did pulse density.

**Figure 4.**
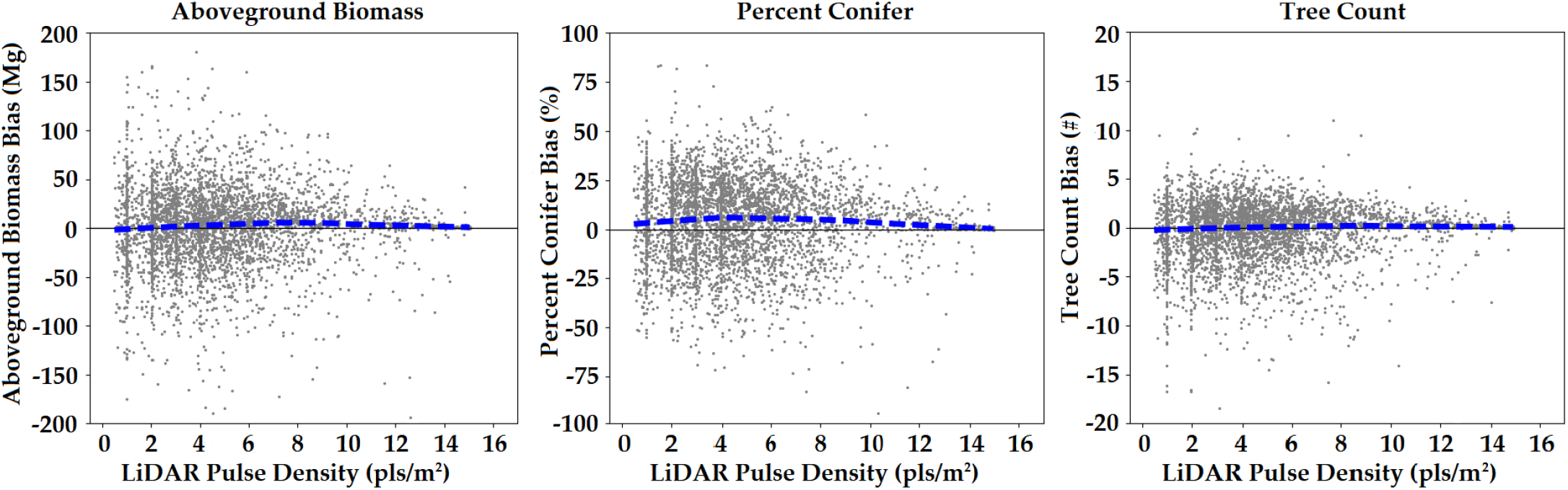
Bias by pulse density in Northern New England using FIA plot-level validation. The dashed blue line is a lowess regression fit of the data.

### 3.3 County-level comparisons

With the region mapped, we compared country level estimates derived by summing the values of our map with FIA county-level design-based estimates. This data is summarized in Table 3. We chose the 38 counties in Northern New England with complete LiDAR coverage. Initially we used all FIA plots within a county measured within two years of the LiDAR acquisition. We discovered however, that a large number of FIA plots without measured trees on them were located in suburban environments with trees. This resulted in underestimates by the FIA data of each forest attribute, so plots that were denoted as having no trees that fell within semi-forested suburbs were removed after manual aerial photo interpretation.

Examining aboveground biomass, our predictions fell within the 95 % confidence interval of the FIA’s in 31/38 counties, and within 97.5 % confidence interval in 33/38 counties. Across the landscape of these counties the FIA estimated 4 % more biomass then our maps. This is to be expected given that our maps frequently had gaps between LiDAR acquisitions and occasionally had missing LiDAR tiles. The FIA’s lack of urban tree sampling likely also played a large role.

Eight of the counties were classified by the US Census Bureau as having an urban population greater than 50 %. In these urbanized counties FIA estimates were an average of 13 % lower than our own when including empty plots in suburban forested areas, and 11 % greater than our own after they were removed.

In estimating percent conifer, 25/38 of our estimates fell within the 95 % confidence interval of the FIA’s estimate. In agreement with the map of residuals, the percentage of conifers was significantly underestimated in 6/11 counties in Maine, and significantly overestimated in 4/8 counties in Vermont. In 16/19 counties in Massachusetts and New Hampshire, our estimate of percent conifer was within the 95 % confidence interval of the FIA’s. Across the entire landscape the two estimates were within 0.2 % of one another.

County level tree count estimates fell within the FIA’s 95 % confidence interval 30/38 times. Once again, counties in Maine with greater numbers of small trees were more likely to be underestimated. In 5/12 Maine counties, mapped estimates of tree count were significantly lower than the FIA’s estimates. Outside of Maine 24/27 counties had mapped estimates in agreement with FIA estimates. This is likely owing to an overall greater stem density in Maine due to more intensive industrial forestry, and a general proclivity of boreal forests to be denser.

**Table 3.** County-level estimates of total aboveground biomass (BIOAG), percent conifer (PC), and tree count (TC) are compared using the FIA’s design-based sampling and summations of our forest inventory maps. Blue denotes mapped estimates that fell within the 95 % confidence interval of the FIA’s estimate, yellow denotes estimates that fell within 97.5 %, red denotes values that estimates that differed from the FIA.

| State | County | FIA BIOAG<br>(Petagrams) | EFI BIOAG<br>(Petagrams) | FIA PC<br>(%) | EFI PC<br>(%) | FIA TC<br>(Millions) | EFI TC<br>(Millions) |
| --- | --- | --- | --- | --- | --- | --- | --- |
| Maine | Cumberland | 21.47 ± 3.48 | 18.75 | 34.45 ± 5.70 | 26.71 | 106.57 ± 17.47 | 89.79 |
| Maine | Hancock | 29.25 ± 3.54 | 29.57 | 55.63 ± 5.09 | 43.25 | 277.55 ± 32.08 | 244.19 |
| Maine | Kennebec | 17.29 ± 2.95 | 16.72 | 30.04 ± 5.40 | 28.86 | 112.41 ± 19.54 | 92.92 |
| Maine | Knox | 7.52 ± 2.08 | 5.48 | 42.67 ± 9.24 | 30.68 | 52.47 ± 13.05 | 35.11 |
| Maine | Lincoln | 9.06 ± 2.07 | 8.31 | 35.02 ± 8.28 | 32.43 | 55.02 ± 14.33 | 48.59 |
| Maine | Penobscot | 56.87 ± 4.64 | 56.21 | 51.11 ± 3.58 | 43.26 | 585.06 ± 41.93 | 452.14 |
| Maine | Piscataquis | 64.96 ± 5.18 | 61.13 | 50.86 ± 3.49 | 46.27 | 654.97 ± 49.96 | 527.91 |
| Maine | Sagadahoc | 6.70 ± 1.51 | 5.18 | 39.53 ± 12.68 | 33.46 | 39.98 ± 9.97 | 29.71 |
| Maine | Waldo | 15.97 ± 2.32 | 11.29 | 38.70 ± 5.44 | 33.92 | 124.14 ± 18.50 | 71.14 |
| Maine | Washington | 42.80 ± 3.86 | 43.11 | 50.94 ± 3.74 | 46.53 | 422.44 ± 32.49 | 404.26 |
| Maine | York | 25.92 ± 3.32 | 22.34 | 30.78 ± 5.26 | 27.52 | 139.91 ± 16.95 | 105.06 |
| Massachusetts | Barnstable | 4.49 ± 1.54 | 4.43 | 36.53 ± 13.00 | 23.96 | 36.25 ± 12.78 | 41.28 |
| Massachusetts | Berkshire | 34.42 ± 4.43 | 32.59 | 18.73 ± 5.05 | 28.67 | 114.48 ± 15.41 | 126.32 |
| Massachusetts | Bristol | 11.92 ± 2.98 | 11.89 | 15.52 ± 5.34 | 23.90 | 52.97 ± 15.64 | 56.44 |
| Massachusetts | Dukes | 0.89 ± 0.54 | 0.85 | 20.49 ± 20.15 | 18.87 | 6.29 ± 3.02 | 8.73 |
| Massachusetts | Essex | 13.98 ± 2.10 | 11.15 | 23.61 ± 5.15 | 18.41 | 57.68 ± 12.13 | 39.25 |
| Massachusetts | Franklin | 23.98 ± 3.75 | 27.30 | 31.35 ± 5.86 | 29.99 | 86.87 ± 14.18 | 93.13 |
| Massachusetts | Middlesex | 20.76 ± 4.31 | 17.68 | 23.50 ± 8.09 | 21.57 | 60.96 ± 15.27 | 65.82 |
| Massachusetts | Nantucket | 0.34 ± 1.19 | 0.06 | 0.00 | 6.99 | 2.49 ± 8.75 | 0.75 |
| Massachusetts | Norfolk | 9.66 ± 3.07 | 7.91 | 14.92 ± 8.76 | 21.77 | 36.71 ± 11.71 | 33.72 |
| Massachusetts | Plymouth | 13.73 ± 3.66 | 14.43 | 33.96 ± 8.64 | 27.24 | 58.78 ± 14.02 | 67.32 |
| Massachusetts | Suffolk | NO PLOTS | 0.33 | NO PLOTS | 10.73 | NO PLOTS | 2.07 |
| Massachusetts | Worcester | 44.63 ± 5.77 | 41.17 | 22.91 ± 4.75 | 24.02 | 144.91 ± 18.86 | 154.38 |
| New Hampshire | Belknap | 10.47 ± 2.42 | 11.06 | 29.18 ± 8.56 | 25.45 | 50.78 ± 11.55 | 45.40 |
| New Hampshire | Cheshire | 26.48 ± 3.24 | 25.58 | 29.09 ± 5.57 | 31.83 | 105.08 ± 14.76 | 98.29 |
| New Hampshire | Hillsborough | 25.83 ± 4.00 | 23.54 | 28.78 ± 5.45 | 27.27 | 102.79 ± 15.68 | 92.32 |
| New Hampshire | Merrimack | 32.48 ± 4.44 | 24.62 | 31.80 ± 4.83 | 27.67 | 134.76 ± 17.09 | 99.81 |
| New Hampshire | Rockingham | 19.07 ± 3.92 | 17.89 | 32.19 ± 6.27 | 22.27 | 72.74 ± 14.78 | 72.68 |
| New Hampshire | Strafford | 9.39 ± 2.43 | 9.76 | 25.51 ± 8.13 | 27.51 | 38.43 ± 10.20 | 42.43 |
| New Hampshire | Sullivan | 16.11 ± 2.58 | 17.01 | 30.85 ± 7.20 | 33.55 | 74.51 ± 12.52 | 72.30 |
| Vermont | Caledonia | 15.18 ± 3.12 | 14.79 | 35.60 ± 8.20 | 33.14 | 85.38 ± 16.62 | 75.95 |
| Vermont | Essex | 10.58 ± 1.59 | 13.95 | 23.61 ± 5.15 | 39.51 | 76.14 ± 9.19 | 92.98 |
| Vermont | Lamoille | 13.73 ± 2.14 | 12.37 | 23.19 ± 8.16 | 35.57 | 60.14 ± 9.54 | 62.42 |
| Vermont | Orange | 18.18 ± 3.16 | 19.18 | 32.09 ± 8.22 | 30.81 | 78.32 ± 14.20 | 79.24 |
| Vermont | Rutland | 30.77 ± 3.00 | 25.65 | 19.01 ± 4.26 | 26.68 | 116.47 ± 11.76 | 110.83 |
| Vermont | Washington | 18.11 ± 2.92 | 19.55 | 29.39 ± 6.60 | 33.57 | 85.94 ± 13.20 | 90.94 |
| Vermont | Windham | 28.02 ± 3.38 | 27.60 | 26.10 ± 6.23 | 33.09 | 102.42 ± 12.16 | 103.68 |
| Vermont | Windsor | 30.81 ± 4.33 | 30.86 | 19.23 ± 5.52 | 29.96 | 112.53 ± 16.81 | 116.98 |

## 3. Discussion

Our results indicate that 3D convolutional neural networks can be used to estimate forest attributes from disparate LiDAR and satellite data. These models outperformed random forest models which use the standard approach for generating forest inventories from LiDAR. They could also be effectively scaled to make regional high resolution maps/estimates which were often statistically equivalent to traditional forest inventories.

### 4.1 Model comparison

Our first objective was to compare LiDAR derived inventory estimates made using CNNs to estimates made using height metrics and random forest. We assessed this in our first phase of validation, in which several models were developed from training data and assessed using withheld plots. Random forest models trained using traditional height metrics and satellite data nearly always had considerably greater error than the two CNNs we trained. This mirrors the results found by Ayrey and Hayes (2018), in which 3D CNNs of varying architectures often outperformed generalized linear models and random forest. These results indicate that deep learning can be a more effective way of modelling LiDAR than the standard approach of height metrics.

In the estimation of species (percent conifer and percent spruce/fir), random forest did outperform the CNNs. These species breakdowns were more likely to rely on satellite spectral data than LiDAR structural data, and so we speculate that random forest did a better job of making use of the satellite covariates than our CNN. The CNN was initially trained to scan LiDAR voxels, the addition of satellite covariates was made after this training, and in such a ways as to concatenate satellite data onto voxel space. This may have been less than ideal. Zhou and Hauser (2017) outlined several methods for including side-channel data into a CNN, and we tried each of them with mixed results before settling on our concatenation method.

The CNNs also did not outperform the random forest models in terms of bias. Half of the random forest models had a lower absolute bias than the CNNs, indicating that both types of models did approximately as well as one another. We did not observe any notable trends in biases by forest attribute. Overall, we believe that the differences in absolute bias between models was often low enough to have been owing to the random variations of the testing data.

The comparison between the CNNs with and without satellite metrics highlighted that the CNN did benefit from spectral and disturbance information. Error decreased when estimating every forest attribute, and bias nearly always decreased as well. In terms of magnitude this improvement was often small, and our results indicate that even without the satellite metrics, a CNN could be trained which outperforms random forest with height and satellite metrics. We chose not to explore which satellite metrics were most useful, as deep learning models of this size run no risk of overfitting with extraneous inputs. However, such an analysis would be possible through a process similar to random forest’s derived importance, and might be useful in narrowing down necessary remote sensing datasets.

We noted that there were numerous advantages to working with a single deep learning model aside from better performance. Our Inception-V3 model took a considerable amount of data and time to train, however, once trained the model could be quickly be applied to large regional LiDAR datasets. A single model predicting all 14 forest attributes presented less of a data-management challenge than 14 separate models. We also hypothesize that our CNN would be less likely to produce estimates in conflict with one another than 14 separate unconstrained models (e.g. more merchantable volume than total volume). Finally, there is a considerable amount of precedence in the field of deep learning for making use of pre-trained model weights to solve novel problems (Pan and Yang 2010, Shin et al. 2016,). The rapid retraining of our CNNs indicates that this model can easily be fine-tuned with local data, retuned with non-local data, or applied to different problems to save modelers the effort of training a large CNN to interpret voxel space with randomized weights. The weights from our CNN could be used to initialize CNNs with other LiDAR-related objectives, such as individual tree crown segmentation or LiDAR classification.

### 4.2 Assessing performance

With the model form decided upon, we mapped all fourteen forest attributes across the study area (Figure 4). We assessed the maps’ performance with our second phase of validation, which made use of independent FIA plots. We assessed accuracy at a supplot-, plot-, and county-level. Our subplot-level error estimates were consistently quite high. The FIA plot locations in this region are subject to considerable error, and the FIA notes that plot location error can be up to 100 m (although in practice most plots are located within 12 m of their measured location, Hoppus and Lister 2007). Examining pixel-level accuracy using these subplots was problematic given the high degree of intra-canopy variation present in 10 m pixels. Our observation that the center subplot (on which the GPS location was taken) resulted in lower map error is a further indication that the locational accuracy of the surrounding subplots is suspect.

Our assessment of plot-level error (the aggregate of the subplots) produced more favorable error results. We achieved nRMSE values of between 40-50 % for those attributes quantifying tree size, which we consider to be a success given the small size of the grid cells used. Estimates of tree count, percent conifer, and deciduous volume we consider to have been made with moderate success with nRMSE values of between 56-61 %. Estimates of *pinus strobus* and spruce/fir species breakdowns we consider to be a failure, with nRMSEs exceeding 100 %. Despite this, these maps may still be of some use to some practitioners when aggregated to a more coarse resolution and binned into categories.

**Figure 4.**
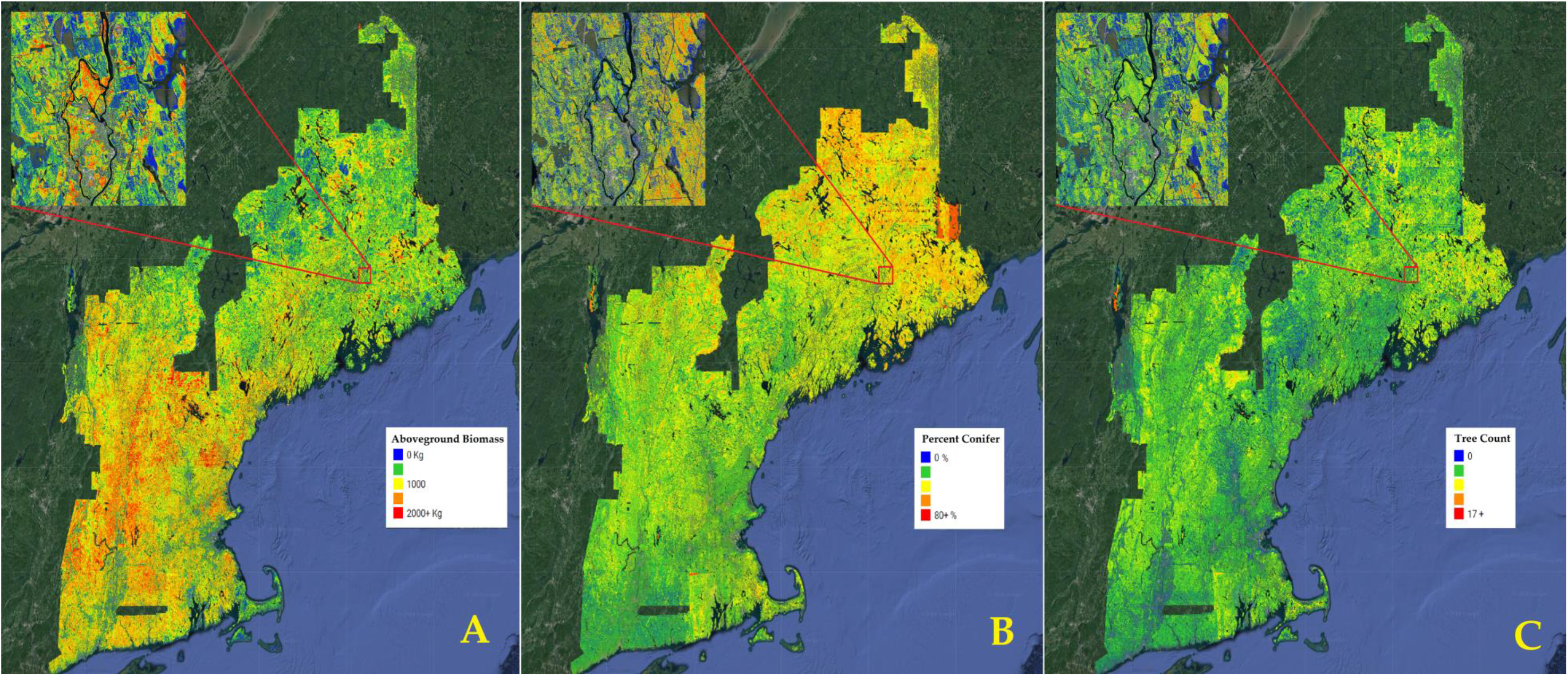
Regional 10 m forest inventory maps developed over New England. (A) Aboveground biomass. (B) Percent conifer. (C) Tree Count.

We assessed model performance at the county-level and in space using aboveground biomass, percent conifer, and tree count. Our map of biomass residuals across Northern New England appeared to be relatively uniform. We noted no major trends in residuals in space, indicating that we are representing biomass across the landscape well. Notably, we did not experience any saturation of high biomass areas as is often the case with regional remote sensing studies (Zhao et al. 2016). Our biomass estimates fell within the 97.5 % confidence of FIA biomass estimates in all but four counties. Those four counties appeared to have little in common in terms of forest characteristics, nearness in space, or population density.

We also noted that in urban and urbanized counties, FIA estimates of biomass could be made to underestimate or overestimate our own, depending on sampling design. Retaining supposedly empty FIA plots placed in suburban areas in which trees were present in aerial imagery resulted in the FIA underestimating biomass relative to our maps. This suggests that our maps are doing a better job of quantifying urban and suburban biomass. By our estimate, this adds up to an additional 13 % biomass in urbanized counties. However, we should stress that none of our training data made use of urban plots, and few of our training plots had trees grown in the open. This improvement may only be the case of some estimate being better than none at all.

The map of percent conifer residuals showed a systematic underestimation of percent conifer in Eastern Maine, and overestimation in Vermont. These areas are inhabited by very high and low proportions of conifers respectively. This trend can also be seen in the county-level comparisons of percent conifer. Examining the predicted versus observed plot confirms that this model underfit the extremes. The map of tree count residuals was similar to the percent conifer in that there was a consistent underestimation of trees in Northern Maine, where stem densities are naturally higher due to the species assemblages and greater industrial disturbance results in younger forests. The predicted versus observed plot highlights a model saturation in very high density forests. Unlike percent conifer, no consistent underestimation of tree count was observed in lower density areas. The county-level estimates parallel this finding, with the majority of the disagreement between the FIA tree count estimates being in Maine in counties with the highest stem densities. Intuitively, one might expect this result given that the structure of very dense forest stands resemble one another despite dramatically different stem densities (e.g. a point cloud representing a stand with 2000 trees per hectare looks very similar to a point cloud representing 2500 trees per hectare). Satellite indices are also sometimes prone to the same type of saturation at very high stem densities (Mohammadi et al. 2010).

Our model’s performance can be summarized as having done a good job at predicting attributes closely related to tree size, having done a moderate job of predicting attributes related to tree density and percent conifer, and having done a poor job of predicting attributes related to species groupings. Other studies modeling forest attribute using LiDAR have likewise had more difficulty in estimating stem density and species (Treitz et al. 2012, Hayashi et al. 2014, Shang et al. 2017, Hudak et al. 2008). We can think of several reasons for why our models have also underperformed here. (1) Although we incorporated satellite spectral indices useful for species estimation, we believe that the model may have not made full use of them. (2) Stem density was often underestimated in high density stands, but qualitatively the maps seemed to suffer from banding in areas with low pulse density LiDAR (<3 pls/m^2^). It is possible that the model was making use of horizontal structural features in the canopy which could not be resolved in low density LIDAR. Ayrey and Hayes (2018) determined that 3D CNNs do make use of horizontal canopy structure (such as the edges of tree crowns). (3) Finally, in re-examining our loss function, we find that 7/14 of our forest attributes were in some way related to tree size, while only one attribute estimated stem density. It is possible that our unweighted loss function inadvertently favored attributes estimating tree size, resulting in a model that identified features in the LiDAR that were more predictive of size.

### 4.4 Mapping errors

Unfortunately, the regional maps of forest attributes suffered from several types of errors. One source of these errors were the LiDAR acquisitions themselves, which often didn’t entirely overlap, or were collected improperly. Missing areas can often be observed between the gaps of the nearly 40 different LiDAR acquisitions over the region. In central Connecticut and Eastern Maine portions of the LiDAR were acquired with improper settings, resulting in vegetation being severely under-represented.

Banding errors occurred with forest attributes that had moderate/worse performance (tree count, percent conifer, and species/volume estimates). These bands were more likely to occur in areas in which pulse density fell below 3 pls/m^2^, and followed scan angle trends. In environments of low pulse density and high scan angle, these attributes were often underestimated, possibly owing to less horizontal structure being captured by the LiDAR. Maps estimating tree size, such as BIOAG, VOL, and BA, had few banding errors.

### 4.5 Our results in context

Several other studies have mapped aboveground biomass in this region and can be used to place our model’s performance in context. In one early example, Kellndorfer et al. used Landsat and radar to map biomass across the Continental United States (CONUS), and achieved RMSE values ranging from 42 – 48 Mg/ha over New England with 30 m pixels (Cartus et al. 2012). In 2008 Blackard et al. used MODIS to predict biomass across the CONUS at a 250 m resolution and achieved an average absolute error in New England ranging between 49.7 – 60.4 Mg/ha. In a more regional study, Cartus et al. (2012) mapped aboveground biomass in the Northeastern United States using radar, and achieved RMSE estimates of between 46 – 47.3 Mg/ha with 150 m pixels, but noted that increasing pixel size dramatically reduced error. A more recent study mapped biomass in New England and Atlantic Canada using Landsat time-series data, this achieved a RMSE of 44.7 Mg/ha using 30 m pixels (Kilbride 2018). In the context of these studies, our aboveground biomass error of 36 Mg/ha at a roughly 20 m resolution (FIA plot-level error), represents a considerable improvement over existing remotely-sensed regional estimates.

Localized studies in experimental forests throughout the region can also be used for comparison. In a 2016 study Hayashi et al. mapped stem volume using LiDAR and achieved RMSEs of 46 m^3^/ha and 82 m^3^/ha in two experimental forests in Maine and New Brunswick (we achieved an error of 62.5 m^3^/ha). In a similar study, Hayashi et al. obtained RMSEs of 4993 trees/ha, 3.68 cm, and 13 m^2^/ha for tree count, QMD, and basal area respectively using 20 m cells at an experimental forest in Maine (Hayashi et al. 2014). Our regional models achieved errors of 293 trees/ha, 4.5 cm, and 7.9 m^2^/ha for these attributes; outperforming the local models in estimating tree count and basal area. Another study at an experimental forest in Massachusetts used large footprint LiDAR and radar to estimate biomass, achieving a RMSE of 66.6 Mg/ha with 25 m cells (Ahmed et al. 2010). Taken collectively, our regional models seem to perform on par or better than local modeling efforts.

### 4.6 Conclusions

In this study we mapped the forests of New England at a 10 m resolution, making estimates of fourteen different forest inventory attributes. We did so through the use of disparate LiDAR datasets as well as satellite spectral, phenological, and disturbance data. Our method of modeling these attributes was somewhat novel, and made use of three dimensional convolutional neural networks which are a form of deep learning. These CNNs outperformed standard modeling techniques, and proved themselves useful for large-scale mapping, making use of disparate data, and increasing data management and computational efficiency.

We validated our maps using an external dataset derived from the USFS’s forest inventory and analysis program. We concluded that the most successful estimates were those attributes that quantified tree size, moderately successful estimates were those that quantified tree density or percent coniferous, and less successful estimates were those that quantified more specific species groupings. In particular, we found our biomass estimates to be in good agreement with the FIA’s across the region.

We believe that both the models and the maps generated by this study will be of use to many other scientists. The weights from the CNN trained here could be used as a starting point to train models making estimates over different forests, or to other LiDAR-related remote sensing problems. Likewise, the maps developed here can assist with wildlife mapping, precision forestry, and carbon stock estimation in the region.

## Acknowledgments

We thank NASA Goddard’s G-LiHT team and NEON for the use of their LiDAR. We thank Kathryn Miller and the National Park Service for the use of their Acadia National Park field data. We thank Eben Sypitowski and Baxter State Park for the use of their Scientific Forest Management Area CFI field plots. We thank Brian Roth and the University of Maine’s Cooperative Forestry Research Unit for the use of their LiDAR and field data, as well as for considerable technical assistance. We thank Keith Kanoti and the University of Maine for the use of their CFI field plots. We thank the New Hampshire Division of Forests and Lands, Caroline A. Fox Research and Demonstration Forest for the use of their CFI field plot data. We thank David Orwig and Harvard Forest for the use of their Megaplot field inventory. We thank Noonan research forest for the use of their stem mapped plots. We thank Laura Kenefic and the U.S. Forest Service’s Penobscot Experimental Forest for the use of their LiDAR and field inventory data.

## Funding details

This work was supported by the Maine Agricultural and Forest Experiment Station, Publication Number XXXX. This project was supported by the USDA National Institute of Food and Agriculture, McIntire-Stennis project number ME0-XXXX through the Maine Agricultural & Forest Experiment Station.

## Conflicts of Interest

The authors declare no conflict of interest.

## Data availability statement

Computer code for this work can be found online at https://github.com/Eayrey/3D-Convolutional-Neural-Networks-with-LiDAR. Generated inventory maps can be found online at XXXXXXXXXXXXXXXXXX

---

**Appendix Table A1.**
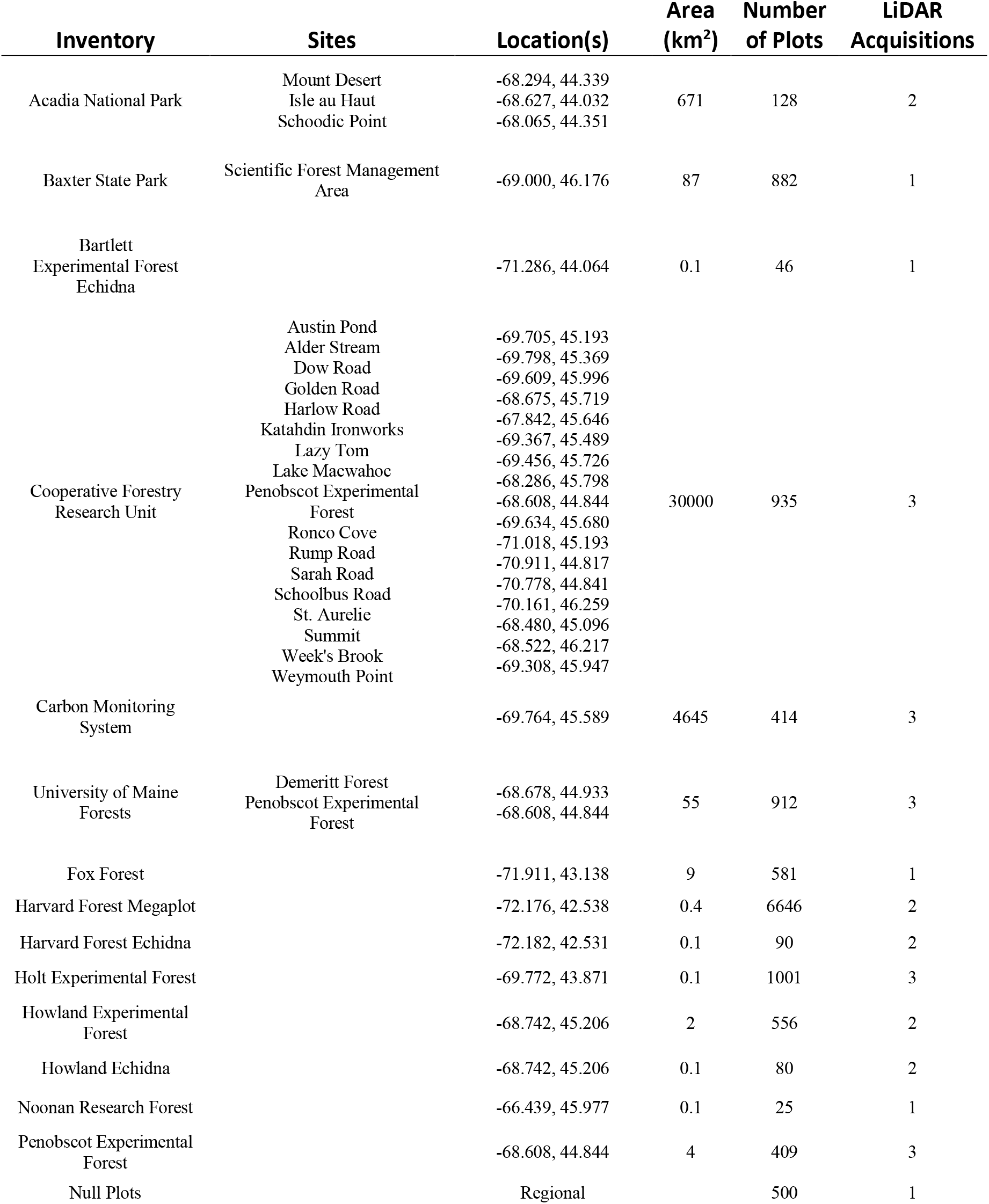
A list of field inventories used for model training and the first phase of validation. Also included, the area those inventories represented, the number of LiDAR field plots, and the number of LiDAR acquisitions. When inventories covered multiple sites, those sites are listed.

## References

Abadi, M., Barham, P., Chen, J., Chen, Z., Davis, A., Dean, J., Devin, M., Ghemawat, S., Irving, G., & Isard, M. (2016). Tensorflow: a system for large-scale machine learning. In, OSDI (pp. 265–283)

Ahmed, R., Siqueira, P., Bergen, K., Chapman, B., & Hensley, S. (2010). A biomass estimate over the harvard forest using field measurements with radar and lidar data. In, Geoscience and Remote Sensing Symposium (IGARSS), 2010 IEEE International (pp. 4768–4771): IEEE

Ayrey, E., & Hayes, D.J. (2018). The Use of Three-Dimensional Convolutional Neural Networks to Interpret LiDAR for Forest Inventory. Remote Sensing, 10, 649

Blackard, J., Finco, M., Helmer, E., Holden, G., Hoppus, M., Jacobs, D., Lister, A., Moisen, G., Nelson, M., & Riemann, R. (2008). Mapping US forest biomass using nationwide forest inventory data and moderate resolution information. Remote Sensing of Environment, 112, 1658–1677

Breiman, L. (2001). Random forests. Machine Learning, 45, 5–32

Cartus, O., Santoro, M., & Kellndorfer, J. (2012). Mapping forest aboveground biomass in the Northeastern United States with ALOS PALSAR dual-polarization L-band. Remote Sensing of Environment, 124, 466–478

Castelluccio, M., Poggi, G., Sansone, C., & Verdoliva, L. (2015). Land use classification in remote sensing images by convolutional neural networks. arXiv preprint arXiv:1508.00092

Dick, A. (2019). Enhanced Forest Inventory (EFI) Adoption in New Brunswick: Progress to Date and Future Directions. In, CIF E-Lecture Series. Canadian Wood Fibre Centre: Natural Resources Canada

García-Feced, C., Tempel, D.J., & Kelly, M. (2011). LiDAR as a tool to characterize wildlife habitat: California spotted owl nesting habitat as an example. Journal of Forestry, 109, 436–443

Genuer, R., Poggi, J.-M., & Tuleau-Malot, C. (2015). VSURF: an R package for variable selection using random forests. The R Journal, 7, 19–33

Ghamisi, P., Höfle, B., & Zhu, X.X. (2017). Hyperspectral and LiDAR data fusion using extinction profiles and deep convolutional neural network. IEEE Journal of Selected Topics in Applied Earth Observations and Remote Sensing, 10, 3011–3024

Goodwin, N.R., Coops, N.C., & Culvenor, D.S. (2006). Assessment of forest structure with airborne LiDAR and the effects of platform altitude. Remote Sensing of Environment, 103, 140–152

Gorelick, N., Hancher, M., Dixon, M., Ilyushchenko, S., Thau, D., & Moore, R. (2017). Google Earth Engine: Planetary-scale geospatial analysis for everyone. Remote Sensing of Environment, 202, 18–27

Guan, H., Yu, Y., Ji, Z., Li, J., & Zhang, Q. (2015). Deep learning-based tree classification using mobile LiDAR data. Remote Sensing Letters, 6, 864–873

Guo, X., Coops, N.C., Tompalski, P., Nielsen, S.E., Bater, C.W., & Stadt, J.J. (2017). Regional mapping of vegetation structure for biodiversity monitoring using airborne lidar data. Ecological informatics, 38, 50–61

Hayashi, R., Kershaw, J.A., & Weiskittel, A. (2015). Evaluation of alternative methods for using LiDAR to predict aboveground biomass in mixed species and structurally complex forests in northeastern North America. MCFNS, 7, 49–65

Hayashi, R., Weiskittel, A., & Kershaw Jr, J.A. (2016). Influence of prediction cell size on LiDAR-derived area-based estimates of total volume in mixed-species and multicohort forests in northeastern North America. Canadian Journal of Remote Sensing, 42, 473–488

Hayashi, R., Weiskittel, A., & Sader, S. (2014). Assessing the feasibility of low-density LiDAR for stand inventory attribute predictions in complex and managed forests of northern Maine, USA. Forests, 5, 363–383

He, K., Zhang, X., Ren, S., & Sun, J. (2016). Deep residual learning for image recognition. In, Proceedings of the IEEE conference on computer vision and pattern recognition (pp. 770–778)

Hoppus, M., & Lister, A. (2007). The status of accurately locating forest inventory and analysis plots using the Global Positioning System. In, In: McRoberts, Ronald E.; Reams, Gregory A.; Van Deusen, Paul C.; McWilliams, William H., eds. Proceedings of the seventh annual forest inventory and analysis symposium; October 3-6, 2005; Portland, ME. Gen. Tech. Rep. WO-77. Washington, DC: US Department of Agriculture, Forest Service: 179–184.

Hudak, A.T., Crookston, N.L., Evans, J.S., Hall, D.E., & Falkowski, M.J. (2008). Nearest neighbor imputation of species-level, plot-scale forest structure attributes from LiDAR data. Remote Sensing of Environment, 112, 2232–2245

Jensen, J.L.R., Humes, K.S., Conner, T., Williams, C.J., & DeGroot, J. (2006). Estimation of biophysical characteristics for highly variable mixed-conifer stands using small-footprint lidar. Canadian Journal of Forest Research, 36, 1129–1138

Junttila, V., Kauranne, T., Finley, A.O., & Bradford, J.B. (2015). Linear Models for Airborne-Laser-Scanning-Based Operational Forest Inventory With Small Field Sample Size and Highly Correlated LiDAR Data. IEEE Transactions on Geoscience and Remote Sensing, 53, 5600–5612

Kangas, A., Astrup, R., Breidenbach, J., Fridman, J., Gobakken, T., Korhonen, K.T., Maltamo, M., Nilsson, M., Nord-Larsen, T., & Næsset, E. (2018). Remote sensing and forest inventories in Nordic countries–roadmap for the future. Scandinavian Journal of Forest Research, 33, 397–412

Kennedy, R.E., Yang, Z., & Cohen, W.B. (2010). Detecting trends in forest disturbance and recovery using yearly Landsat time series: 1. LandTrendr—Temporal segmentation algorithms. Remote Sensing of Environment, 114, 2897–2910

Kilbride, J.B. (2018). Forest Disturbance Detection and Aboveground Biomass Modeling Using Moderate-Resolution, Time-Series Satellite Imagery

Ko, C., Kang, J., & Sohn, G. (2018). Deep Multi-task Learning for Tree Genera Classification. ISPRS Ann. Photogramm. Remote Sens. Spat. Inf. Sci, 153–159

Krizhevsky, A., Sutskever, I., & Hinton, G.E. (2012). Imagenet classification with deep convolutional neural networks. In, Advances in neural information processing systems (pp. 1097–1105)

LeCun, Y., & Bengio, Y. (1995). Convolutional networks for images, speech, and time series. The handbook of brain theory and neural networks, 3361, 1995

LeCun, Y., Bengio, Y., & Hinton, G. (2015). Deep learning. nature, 521, 436

Li, R., Weiskittel, A., Dick, A.R., Kershaw Jr, J.A., & Seymour, R.S. (2012). Regional stem taper equations for eleven conifer species in the Acadian region of North America: development and assessment. Northern Journal of Applied Forestry, 29, 5–14

Li, W., Fu, H., Yu, L., & Cracknell, A. (2016). Deep learning based oil palm tree detection and counting for high-resolution remote sensing images. Remote Sensing, 9, 22

Lim, K.S., & Treitz, P.M. (2004). Estimation of above ground forest biomass from airborne discrete return laser scanner data using canopy-based quantile estimators. Scandinavian Journal of Forest Research, 19, 558–570

Maturana, D., & Scherer, S. (2015). 3d convolutional neural networks for landing zone detection from lidar. In, Robotics and Automation (ICRA), 2015 IEEE International Conference on (pp. 3471–3478): IEEE

McGaughey, R.J. (2009). FUSION/LDV: Software for LIDAR data analysis and visualization. US Department of Agriculture, Forest Service, Pacific Northwest Research Station: Seattle, WA, USA, 123

Means, J.E., Acker, S.A., Fitt, B.J., Renslow, M., Emerson, L., & Hendrix, C.J. (2000). Predicting forest stand characteristics with airborne scanning lidar. Photogrammetric Engineering and Remote Sensing, 66, 1367–1372

Mohammadi, J., Shataee Joibary, S., Yaghmaee, F., & Mahiny, A. (2010). Modelling forest stand volume and tree density using Landsat ETM+ data. International Journal of Remote Sensing, 31, 2959–2975

Næsset, E. (2005). Assessing sensor effects and effects of leaf-off and leaf-on canopy conditions on biophysical stand properties derived from small-footprint airborne laser data. Remote Sensing of Environment, 98, 356–370

Næsset, E. (2009). Effects of different sensors, flying altitudes, and pulse repetition frequencies on forest canopy metrics and biophysical stand properties derived from small-footprint airborne laser data. Remote Sensing of Environment, 113, 148–159

Naesset, E. (1997). Determination of mean tree height of forest stands using airborne laser scanner data. ISPRS Journal of Photogrammetry and Remote Sensing, 52, 49–56

Nilsson, M., Nordkvist, K., Jonzén, J., Lindgren, N., Axensten, P., Wallerman, J., Egberth, M., Larsson, S., Nilsson, L., & Eriksson, J. (2017). A nationwide forest attribute map of Sweden predicted using airborne laser scanning data and field data from the National Forest Inventory. Remote Sensing of Environment, 194, 447–454

Ørka, H.O., Næsset, E., & Bollandsås, O.M. (2010). Effects of different sensors and leaf-on and leaf-off canopy conditions on echo distributions and individual tree properties derived from airborne laser scanning. Remote Sensing of Environment, 114, 1445–1461

Pan, S.J., & Yang, Q. (2010). A survey on transfer learning. IEEE Transactions on knowledge and data engineering, 22, 1345–1359

Patenaude, G., Hill, R., Milne, R., Gaveau, D., Briggs, B., & Dawson, T. (2004). Quantifying forest above ground carbon content using LiDAR remote sensing. Remote Sensing of Environment, 93, 368–380

Pflugmacher, D., Cohen, W.B., & Kennedy, R.E. (2012). Using Landsat-derived disturbance history (1972–2010) to predict current forest structure. Remote Sensing of Environment, 122, 146–165

Qi, C.R., Su, H., Mo, K., & Guibas, L.J. (2017). Pointnet: Deep learning on point sets for 3d classification and segmentation. Proc. Computer Vision and Pattern Recognition (CVPR), IEEE, 1, 4

Rizaldy, A., Persello, C., Gevaert, C., Oude Elberink, S., & Vosselman, G. (2018). Ground and Multi-Class Classification of Airborne Laser Scanner Point Clouds Using Fully Convolutional Networks. Remote Sensing, 10, 1723

Russakovsky, O., Deng, J., Su, H., Krause, J., Satheesh, S., Ma, S., Huang, Z., Karpathy, A., Khosla, A., & Bernstein, M. (2015). Imagenet large scale visual recognition challenge. International Journal of Computer Vision, 115, 211–252

Russell, M.B., & Weiskittel, A.R. (2011). Maximum and largest crown width equations for 15 tree species in Maine. Northern Journal of Applied Forestry, 28, 84–91

Shang, C., Treitz, P., Caspersen, J., & Jones, T. (2017). Estimating stem diameter distributions in a management context for a tolerant hardwood forest using ALS height and intensity data. Canadian Journal of Remote Sensing, 43, 79–94

Shin, H.-C., Roth, H.R., Gao, M., Lu, L., Xu, Z., Nogues, I., Yao, J., Mollura, D., & Summers, R.M. (2016). Deep convolutional neural networks for computer-aided detection: CNN architectures, dataset characteristics and transfer learning. IEEE transactions on medical imaging, 35, 1285–1298

Silva, C.A., Crookston, N.L., Hudak, A.T., Vierling, L.A., Klauberg, C., & Silva, M.C.A. (2017). Package ‘rLiDAR’. In

Szegedy, C., Liu, W., Jia, Y., Sermanet, P., Reed, S., Anguelov, D., Erhan, D., Vanhoucke, V., & Rabinovich, A. (2015). Going deeper with convolutions. In, Proceedings of the IEEE conference on computer vision and pattern recognition (pp. 1–9)

Szegedy, C., Vanhoucke, V., Ioffe, S., Shlens, J., & Wojna, Z. (2016). Rethinking the inception architecture for computer vision. In, Proceedings of the IEEE conference on computer vision and pattern recognition (pp. 2818–2826)

Taigman, Y., Yang, M., Ranzato, M.A., & Wolf, L. (2014). Deepface: Closing the gap to human-level performance in face verification. In, Proceedings of the IEEE conference on computer vision and pattern recognition (pp. 1701–1708)

Team, B.M. (2018). Microsoft Releases 125 million Building Footprints in the US as Open Data. In, Bing blogs: Microsoft

Treitz, P., Lim, K., Woods, M., Pitt, D., Nesbitt, D., & Etheridge, D. (2012). LiDAR sampling density for forest resource inventories in Ontario, Canada. Remote Sensing, 4, 830–848

Villikka, M., Packalén, P., & Maltamo, M. (2012). The suitability of leaf-off airborne laser scanning data in an area-based forest inventory of coniferous and deciduous trees. Silva Fennica, 46

Weinstein, B., Marconi, S., Bohlman, S., Zare, A., & White, E. (2019). Individual tree-crown detection in RGB imagery using self-supervised deep learning neural networks. bioRxiv, 532952

Weiskittel, A., Russell, M., Wagner, R., & Seymour, R. (2012). Refinement of the Forest Vegetation Simulator Northeast variant growth and yield model: Phase III. Cooperative Forest Research Unit. Orono, ME: University of Maine, School of Forest Resources, 96–104

White, J.C., Arnett, J.T., Wulder, M.A., Tompalski, P., & Coops, N.C. (2015). Evaluating the impact of leaf-on and leaf-off airborne laser scanning data on the estimation of forest inventory attributes with the area-based approach. Canadian Journal of Forest Research, 45, 1498–1513

White, J.C., Wulder, M.A., Varhola, A., Vastaranta, M., Coops, N.C., Cook, B.D., Pitt, D., & Woods, M. (2013). A best practices guide for generating forest inventory attributes from airborne laser scanning data using an area-based approach. The Forestry Chronicle, 89, 722–723

Woodall, C.W., Heath, L.S., Domke, G.M., & Nichols, M.C. (2011). Methods and equations for estimating aboveground volume, biomass, and carbon for trees in the US forest inventory, 2010

Woods, M., Pitt, D., Penner, M., Lim, K., Nesbitt, D., Etheridge, D., & Treitz, P. (2011). Operational implementation of a LiDAR inventory in Boreal Ontario. The Forestry Chronicle, 87, 512–528

Wulder, M.A., Bater, C.W., Coops, N.C., Hilker, T., & White, J.C. (2008). The role of LiDAR in sustainable forest management. The Forestry Chronicle, 84, 807–826

Yi, D., Zhou, M., Chen, Z., & Gevaert, O. (2016). 3-D convolutional neural networks for glioblastoma segmentation. arXiv preprint arXiv:1611.04534

Zhang, X., Friedl, M.A., & Schaaf, C.B. (2006). Global vegetation phenology from Moderate Resolution Imaging Spectroradiometer (MODIS): Evaluation of global patterns and comparison with in situ measurements. Journal of Geophysical Research: Biogeosciences, 111

Zheng, D., Rademacher, J., Chen, J., Crow, T., Bresee, M., Le Moine, J., & Ryu, S.-R. (2004). Estimating aboveground biomass using Landsat 7 ETM+ data across a managed landscape in northern Wisconsin, USA. Remote Sensing of Environment, 93, 402–411

Zhao, P., Lu, D., Wang, G., Wu, C., Huang, Y., & Yu, S. (2016). Examining spectral reflectance saturation in Landsat imagery and corresponding solutions to improve forest aboveground biomass estimation. Remote Sensing, 8, 469.

Zhou, Y., & Hauser, K. (2017). Incorporating side-channel information into convolutional neural networks for robotic tasks. In, Robotics and Automation (ICRA), 2017 IEEE International Conference on (pp. 2177–2183): IEEE

